# Facilitated Dissociation of Nucleoid Associated Proteins from DNA in the Bacterial Confinement

**DOI:** 10.1101/2021.03.11.434965

**Authors:** Zafer Koşar, A. Göktuĝ Attar, Aykut Erbaş

**Affiliations:** UNAM-National Nanotechnology Research Center and Institute of Materials Science & Nanotechnology, Bilkent University, Ankara 06800, Turkey; Department of Molecular Biology & Genetics, Bilkent University, Ankara 06800, Turkey

## Abstract

Transcription machinery depends on the temporal formation of protein-DNA complexes. Recent experiments demonstrated that lifetime of the complex can also affect transcription. In parallel, *in vitro* single-molecule studies showed that nucleoid-associated proteins (NAPs) leave the DNA rapidly as the bulk concentration of the protein increases via facilitated dissociation (FD). Nevertheless, whether such concentration-dependent mechanism is functional in a bacterial cell, in which NAP levels and the 3D chromosomal structure are often coupled, is not clear *a priori*. Here, by using extensive coarse-grained molecular simulations, we model the unbinding of specific and nonspecific dimeric NAPs from a high-molecular-weight circular DNA molecule in a cylindrical structure mimicking the cellular confinement of a bacterial chromosome. Our simulations show that physiologically relevant peak protein levels (tens of micromolar) lead to highly compact chromosomal structures. This compaction results in rapid off rates (shorter DNA-residence times) but only for specifically DNA-binding NAPs such as the factor for inversion stimulation (Fis). Contrarily, for nonspecific NAPs, the off rates decrease as the protein levels increase, suggesting an inverse FD pattern. The simulations with restrained chromosome models reveal that this inverse response is due to DNA-segmental fluctuations, and that chromosomal compaction is in favor of faster protein dissociation. Overall, our results indicate that cellular-concentration level of a structural DNA-binding protein can be highly intermingled with its DNA-residence time.

## I. INTRODUCTION

Bacterial transcription factors (TFs) are DNA-binding proteins that can dynamically regulate RNA polymerase’s involvement in transcription by reversibly binding to DNA [1, 2]. Conventionally, binding of a TF to its DNA target site is considered sufficient for the functionality of the corresponding TF (i.e., activation or repression of the target gene). However, a growing body of work has demonstrated that the time that a TF spends along the genome (i.e., DNA-residence time) could contribute to regulation of TF functions [3, 4]. Furthermore, this effect is more pronounced if the TF has diverse transcriptional functionality [5].

Recently, a concentration-dependent mechanism referred to as facilitated dissociation (FD) has been discovered to alter the residence times of key bacterial TFs along with a wide array of DNA-binding proteins [6– 17]. The molecular mechanism of FD is explained by the competition between DNA-bound proteins and solution-phase binding competitors for the same binding site [18, 19]. The competition can be either for sites on the protein (if the competitor is DNA) or sites on the DNA (if the competitor is protein) [20]. Importantly, FD leads to shorter residence times (higher off-rates) as the concentration of solution-phase competitors is increased. Remarkably, among the DNA-binding proteins exhibiting the FD type of dissociation *in vitro* are major nucleoid-associated proteins (NAPs) such as the factor for inversion stimulation (Fis) and HU with diverse transcriptional and structural roles [12, 21]. In single-molecule studies, DNA-residence times of the proteins dropped to minutes from hours in response to concentration increases on the order of several hundred nanomolar [6, 10, 11]. At the cellular level, major NAPs can vary their concentrations even more drastically [22–24]; Fis can increase its cellular concentration reversibly from vanishing levels to 60.000 copies per cell (∼ 100 *µ*M after assuming a cytoplasmic volume of 1*µ*m^3^) in the exponential phase [23, 25]. Nonspecific NAPs such as HU or H-NS also undergo large concentration fluctuations at different stages of the bacterial growth cycle but their levels do not drop below ∼ 10 micromolar ranges [22, 26]. In principle, such concentration fluctuations should easily result in an FD-like mechanism. Nevertheless, structural NAPs can also alter the 3D organization of chromosome by forming dense multiprotein-DNA complexes in a concentration-dependent manner [27–30]. Consequently, elevated protein levels can change the very environment, from which the proteins dissociate and put the functionality of FD at the cellular level into question.

Consistently, when Fis molecules dissociated from either sparsely surface-grafted DNA binding sites, or extended DNA chains or chromosomes, they exhibited faster off-rates with increasing protein concentration [6, 10, 11]. However, when Fis molecules took part in the coating and looping of DNA molecules, they were shown to form highly-stable DNA protein-complexes (i.e., the DNA-residence time is higher than the experimental time window [28]). In separate experiments, when competing Fis were replaced by nucleic acid segments, residence times were also shown to decrease via a DNA-segmental type of FD [20], suggesting a potential relation between chromosome compaction and dissociation kinetics. In accord, several pioneering particle-tracking studies on metaloregulator CueR and ZuR provided evidence that DNA-residence times change with chromosome organization in live cells. However, cellular abundance of those proteins is only on the order of ∼ 1 *µ*M [31, 32], thus, they cannot alter the chromosome structure unlike the structural NAPs. Apart from specifically interacting Fis, nonspecifically interacting HU also exhibited a concentration-dependent DNA binding stability, which was shown to be highly sensitive to DNA substrate conformation and competitor DNA fragments in solution [29]. Another nonspecific NAP, H-NS, also exhibited a dissociation mechanism strongly depending on the type of protein-DNA complexation that it forms [33]. Altogether, the experimental evidence suggests that organizational changes in chromosomal structure resulting from DNA-protein complexation can affect the dissociation kinetics of the proteins. However, how the concentration variations of the proteins contribute to this effect is not clear and demands a systematic study. Such resolving would require focusing on specific and nonspecific proteins individually, at least in a native-like environment, which may be achieved via well-designed *in silico* methods.

Here, we aim to demonstrate a latent regulation mechanism involving cellular NAP levels and higher order chromosome organization by addressing the functionality of FD at the cellular level. Specifically, we focus on the interplay between NAP concentration, protein off-rates, and protein-induced chromosome organization in a model cellular confinement by using a large-scale and coarse-grained molecular dynamics (MD) model of a *E. coli* bacterium (Figure 1). Similar models have been used in the previous studies on bacterial chromosome organization [34–36] but with no attention to the protein dissociation kinetics.

**FIG. 1.**
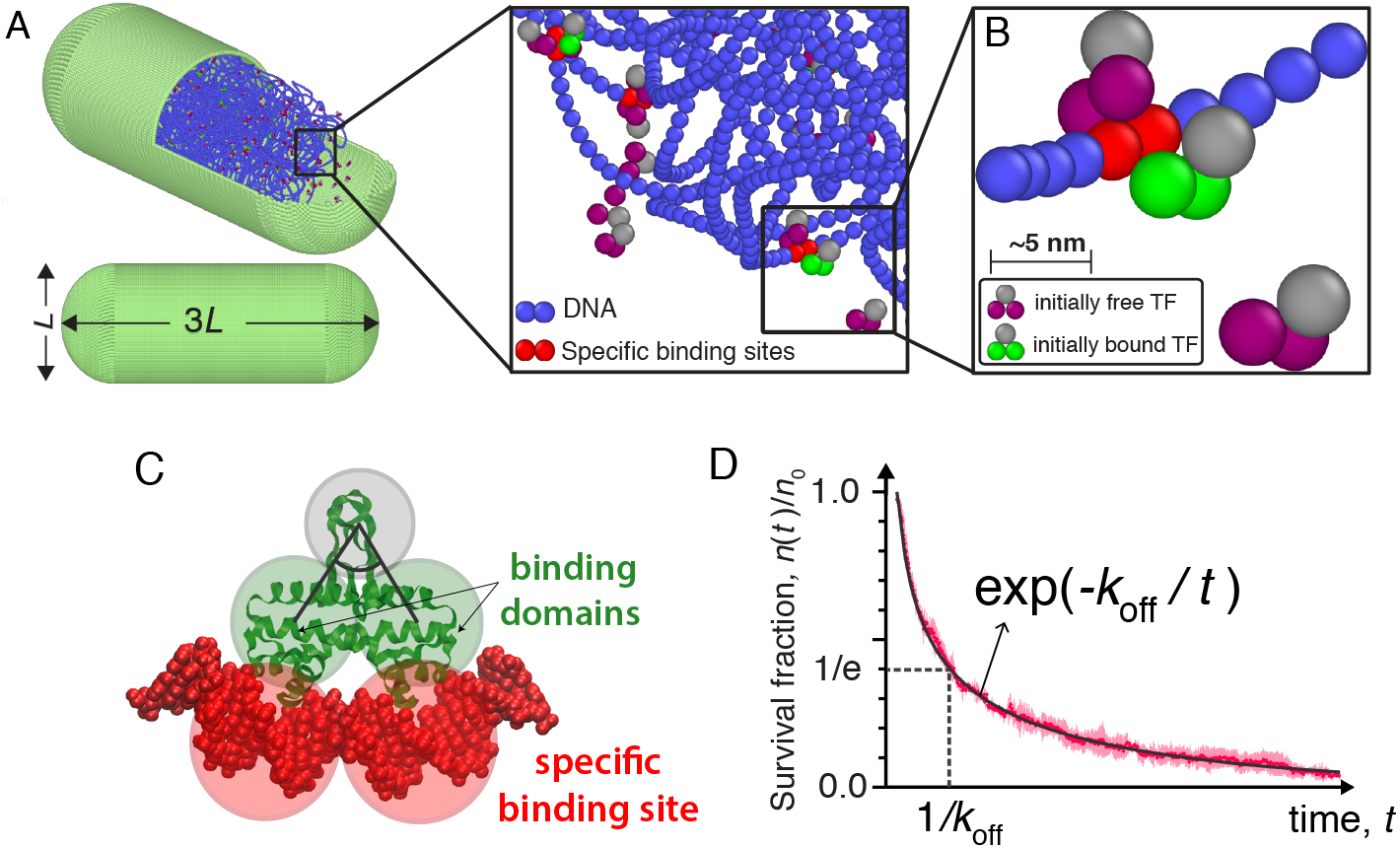
The schematics of the molecular dynamics simulation model. A) The DNA polymer is confined inside a rigid concentric shell (light green beads) together with initially (i.e., at *t* = 0) bound and free proteins. The scale *L* corresponds to 200 nm. B) Closer look into a single specific binding site (red beads) surrounded by nonspecific DNA (blue beads), with which bound and free dimeric proteins can dynamically bind to and unbind from. C) Schematics of the coarse-grained protein model based on the dimeric Fis protein, where two binding domains (green beads) are joined by a hinge bead (gray). D) Extraction of the off-rate, *k*_off_, from the time-dependent decay of the normalized number fraction of DNA-bound proteins (i.e., survival fraction), *n*(*t*)*/n*_0_ (see Eq. 1).

In our simulations, more than one hundred specific (e.g., Fis) or nonspecific (e.g., HU) NAPs with dimeric nature are allowed to fall off simultaneously from their binding sites in the presence of up to several thousand copies of solution-phase proteins. The proteins can dynamically interact (i.e., unbind and rebind) with a high-molecular weight self-avoiding DNA polymer and change the conformation of the polymer dynamically. This setup allows us to precisely track all the proteins in the 3D cell environment simultaneously without any need for molecular labels while monitoring their cumulative effect on the higher-order chromosomal structure. Thus, our computational experiments do not suffer from *in vivo* experimental limitations such as proteins escaping from the 2D-measurement volume or short bleaching times of fluorescence labels.

Interrogation of various protein concentrations and DNA-protein attractions reveal that FD could occur in the cellular confinement, particularly when the chromosome has an open structure. At several tens of micromolar concentrations, the chromosome is collapsed by well-ordered protein-DNA complexes (e.g., doughnut-shaped protein clusters, filamentous structures, and protein networks). Consequently, NAPs exhibit a rich behaviour of dissociation kinetics depending on the sequence-specificity of the proteins.

## II. MATERIALS AND METHODS

### A. Molecular model of bacterial nucleoid

In MD simulations, we modelled bacterial DNA as a single circular polymer by using the coarse-grained Kremer-Grest (KG) self-avoiding bead-spring chain in implicit solvent (Figure 1) [37, 38]. All steric and bond potentials are defined in the model (see Supplementary Data for details). The nucleic acid chain has 1.2 × 10^5^ base pairs (bp), which is modelled by *N* = 12000 KG beads. Each bead represents ∼ 10 bp (i.e., 3.4 nm). The proteins are modelled by three KG beads based on the crystal structure of the DNA-Fis complex (Figure 1B,C) [39]. In the model, two DNA-binding domains are joined by a hinge, which provides a harmonic angle potential to penalize the deviations from the average “cherry” shape of the dimeric protein. The hinge does not bear any attraction towards DNA or other proteins. The semiflexible DNA (persistence length of ∼ 150 bp or ∼ 50 nm) and proteins are confined by a rigid concentric cylinder (also composed of beads) with an aspect ratio of 3 (Figure 1A). The volume of the confinement (1.7 × 10^−2^*µ*m^3^) is adjusted in accordance with the volume fraction of the nucleic acids in *E. coli* (i.e., ∼ 1%). The initial pervaded volume of DNA polymer is set to occupy ∼ 40% of the cell volume to mimic the nucleiod structure [40].

The total number of binding sites along the DNA chain is *n*_0_ = 120. This number is lower than the consensus sequences determined [41] but high enough to perform statistical analyses for unbinding kinetics [42]. At the beginning of a simulation (i.e., *t* = 0), each specific binding site, which is 1000 bp apart from the nearest neighbouring sites, is occupied by a single protein. To monitor the concentration-dependent dissociation, a prescribed concentration of initially free proteins are added at random positions in the simulation box in addition to the *n*_0_ initially DNA-bound proteins. The total protein concentration in cells ranges between ∼ 10 *µ*M (only initially DNA-bound proteins) and 210 *µ*M, which correspond to 120 and 2225 protein copies per cell, respectively.

A DNA-binding protein binds to DNA substrate via weak molecular interactions. Typically, the attraction between DNA and a small TF protein such as Fis is estimated to be around 20-30*k*_B_*T* (i.e., 13-20 kcal/mol) [28, 43, 44], which can lead to hours-long dissociation times for an isolated protein-DNA complex [6, 10]. To observe the dissociation of the DNA-bound proteins within the practically achievable simulation times, we reduced the DNA-protein specific binding attractions and tested that the energy re-scaling only reduces dissociation times without changing the underlying physical mechanisms in accord with the universal behaviour of FD [45] (see Supplementary Data and Supplementary Figures S1 and S2).

All simulations were run on Lammps MD package [46] on parallel 40-core processors. Error bars are calculated by averaging the results of three independent runs and not shown if they are smaller than the corresponding symbol size. Python NumPy package [47] and VMD or OVITO [48, 49] are used for data analysis and the visualizations, respectively

### B. Calculation of off rates

As time progresses in the simulations, the *n*_0_ initially DNA-bound proteins dissociate into the cellular volume from their binding sites. The number of proteins remaining DNA-bound, *n*(*t*), is monitored as a function of the simulation time, *t*. If a bound protein diffuses out of a spherical region with a radius *R*_*c*_ = 8.5 nm centred around the centre of mass of the binding site (red beads in Figure 1), the protein is tagged as dissociated. The distance *R*_*c*_ roughly corresponds to the self-diffusion distance of the protein. If a dissociated protein re-binds to DNA, it is not counted in the surviving bound proteins, *n*(*t*). The survival-fraction data is fit by a single exponential to determine the off-rates,

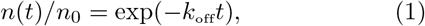

where *k*_off_ is the off-rate with the units of simulation time, which is matched to the self-diffusion time of a protein with size 6.8 nm (∼250 ns) [34]. A sample decay curve is shown in Figure 1D. Further details on the MD simulations and estimations are provided in the Supplementary Data.

## III. RESULTS

### A. The dissociation of specific DNA-binding proteins shows the characteristics of facilitated dissociation

In order to investigate the FD of specific proteins in the confinement of a prokaryotic cell, we monitored the initial dissociation of DNA-bound proteins off their specific binding sites in the absence or presence of prescribed concentrations of initially unbound proteins (Figure 2). Once the proteins dissociate, they can rebind to DNA polymer at the same or another specific site. Alternatively, the protein can bind to DNA nonspecifically but with a lower affinity or stay free inside the cellular volume [21].

**FIG. 2.**
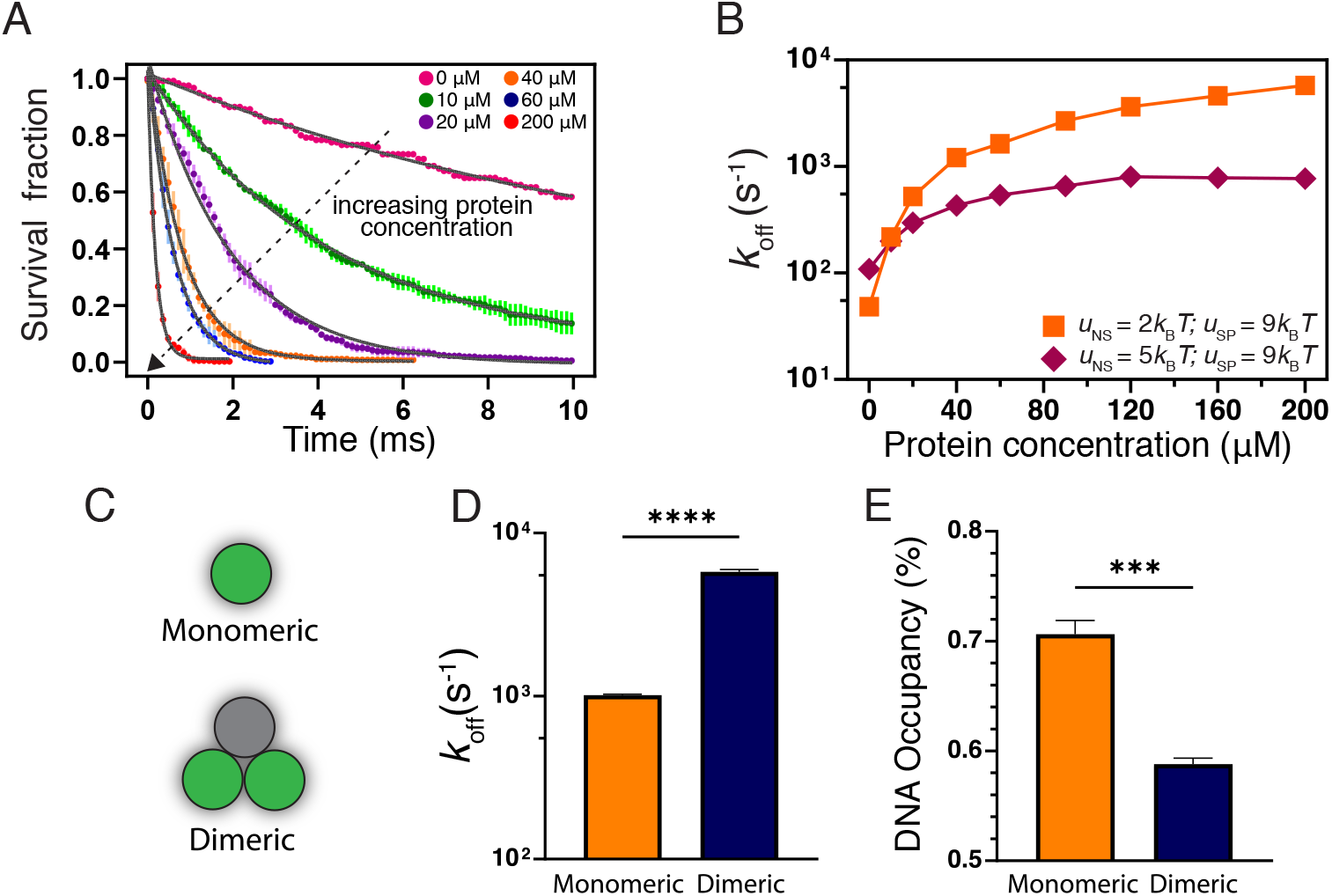
The evidence for the facilitated dissociation in the cellular confinement. A) Representative rescaled survival fraction data of *n*_0_ = 120 initially DNA-bound proteins for various concentrations of initially unbound proteins. The proteins interact with DNA via specific and nonspecific attractions of *u*_SP_ = 9*k*_B_*T* and *u*_NS_ = 2*k*_B_*T*, respectively. The solid curves are the exponential fits (Eq. 1). The regions with lighter colors indicate the error bars. B) The off-rates extracted from the survival-fraction data as a function of unbound protein concentration for various protein-DNA attractions. The data points are joined to guide the eye. C) Monomeric and Dimeric TF models. D) The comparison of the off-rates and E) DNA occupancy of the proteins obtained from the separate simulations with monomeric and dimeric protein models in the presence of initially 200 *µ*M unbound proteins with corresponding valency.

In Figure 2, we show representative dissociation curves of initially DNA-bound proteins leaving their specific binding sites to be replaced by solution-phase proteins at various concentrations. In the case of spontaneous dissociation (SD), for which bound proteins fall off from their specific binding sites in the absence of solution-phase proteins, the dissociation of DNA-bound proteins is significantly slow (i.e., less than 100% of the proteins leave their binding sites within the maximum simulation time) (top curve in Figure 2A). However, as the concentration of initially unbound proteins is increased, the dissociation curves decay more rapidly, indicating that DNA-bound proteins stay on average for a shorter time on their specific binding sites (Figure 2A). For instance, at a protein concentration of 60 *µ*M (1 protein copy per 160 bp), almost all initially DNA-bound proteins leave their specific binding sites after several milliseconds (blue curve in Figure 2A). This concentration-dependent behaviour of dissociation curves is robust and observed for higher DNA-protein attractions (Figure 2B and Supplementary Figure S2) and, consistent with the previous dissociation measurements in the single-molecule experiments [6, 10].

Next, to observe if the dissociation responses follow a typical FD pattern, we extract the off-rates by fitting each dissociation curve by a single exponential function (i.e., Eq. 1). As the binding attraction is increased or the concentration is decreased, the extraction of the off-rates is rather limited due to the slow dissociation of DNA-bound proteins (Figure 2A and Supplementary Figure S2). Hence, in our off-rate calculations, we only use a dissociation curve if more than 40% of DNA-bound proteins dissociate in the corresponding simulation. As expected from the conventional FD behaviour [6, 10], the calculated off-rates of the specifically binding proteins exhibit a consistent increase with increasing concentration of initially free proteins (Figure 2B). Over a concentration range of 10 − 200 *µ*M, the off rates accelerate up to 10-fold as compared to their SD values.

The overall profile of the off rates in Figure 2B is also consistent with the theoretical model [10, 18]: As the concentration increase, a linear increase in the off rates at the low concentrations (taking place within a concentration range of ∼ 10 − 50 *µ*M) is followed by a much weaker increase (Figure 2B). Notably, as the nonspecific attraction between the proteins and DNA becomes weaker (i.e., the value of *u*_NS_ is dropped from 5*k*_B_*T* to 2*k*_B_*T*), the effect of the concentration rise on the off rates becomes stronger (squares versus diamonds in Figure 2B). We will further discuss the effect of nonspecific interactions on dissociation kinetics in the next sections. While the observed concentration-dependence of the off rates qualitatively agrees well with the FD mechanism [6, 10, 21], the off-rate values in the simulations are higher than those observed experimentally, which we attribute to the rescaling of the protein-DNA attraction energies. Nevertheless, the systematic increases in the off rates within the studied concentration range demonstrates the drastic effect of solvated proteins on the dissociation kinetics of specifically DNA-binding proteins in bacterial nucleoid.

### B. Monomeric proteins have a weaker dissociation response than dimeric proteins

Another hallmark of FD is its relation to the multivalency of the DNA-binding proteins [18]. The multivalency can allow proteins to visit microscopically dissociated states on DNA (e.g., a dimeric protein may bind to DNA via one of its binding domains), and thus, to expose the binding site for the invasion of solution-phase proteins[10, 18, 19]. In our default dimeric protein model, the multivalency is achieved by the two independent binding domains (Figure 1C). To test if the concentration-dependent off rates that we observe are related to FD, we decide to replace all the dimeric proteins with their monomeric counterparts in a subset of test simulations. Since monomeric proteins are composed of single beads, they can be either in DNA-bound or free states (Figure 2C). Hence, monomeric proteins should exhibit either none or very weak FD response. In Figure 2D, we compare the off rates of monomeric and dimeric proteins when there are equal amount of unbound proteins in the cell with corresponding valency. Although both protein types bind to the DNA with the same attraction (i.e., specific *u*_SP_ = 9*k*_B_*T* and nonspecific *u*_NS_ = 2*k*_B_*T*) and exhibit similar SD behaviour (i.e., *k*_off_ *<* 10^−5^ *s*^−1^), the off rate of dimeric proteins is roughly one order of magnitude faster than that of monomeric proteins at the same protein level (Figure 2D). Consistently, on average more monomeric proteins occupy the DNA as compared to the dimeric proteins (Figure 2E). This behaviour suggests that the phenomenon that we observe is indeed FD, and it weakens as the multivalency of the proteins is decreased.

### C. Nonspecific attraction between DNA polymer and proteins affects the facilitated dissociation

The specifically interacting NAPs such as Fis can exhibit finite and measurable dissociation constants towards a large set of nonspecific sequences with diverse binding affinities [50]. Further, NAPs such HU or H-NS bind to DNA sequence nonspecifically, and in principle, their binding sites are surrounded by binding sites with equal affinity [22]. Hence, to systematically investigate the role of DNA-protein nonspecific interactions on the FD mechanism, we run simulations for various nonspecific attractions ranging between 0 ≤ *u*_NS_ ≤ *u*_SP_ by fixing the strength of specific DNA-protein attraction, *u*_SP_. In doing so, we can model specifically interacting NAPs (i.e., *u*_NS_ *< u*_SP_) as well as proteins interacting with DNA sequence-independently (i.e., *u*_NS_ = *u*_SP_).

Our simulations reveal the distinct dissociation responses of specific and nonspecific DNA-binding proteins to the increasing free protein levels. Figure 3A shows the off rates for two biologically relevant concentrations, 20 and 60 *µ*M (i.e., 1 protein per roughly 400 and 160 bp, respectively), as a function of the nonspecific attraction strength, *u*_NS_. If the proteins interact specifically with DNA, their off rates increase as a smooth saturation function with increasing protein levels and undergo an apparent FD (Figures 2A and 3). That is, the higher the protein level is, the quicker the DNA-bound protein dissociates. However, as the nonspecific attraction between DNA and proteins is increased, the off-rate becomes visibly independent of the protein concentration at a nonspecific attraction value of around 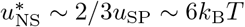 (Figure 3A). We observe this behaviour also for higher values of the specific attraction (Supplementary Figure S3), suggesting that this phenomenon is not a side effect of the lower binding energies that we use in the simulations.

**FIG. 3.**
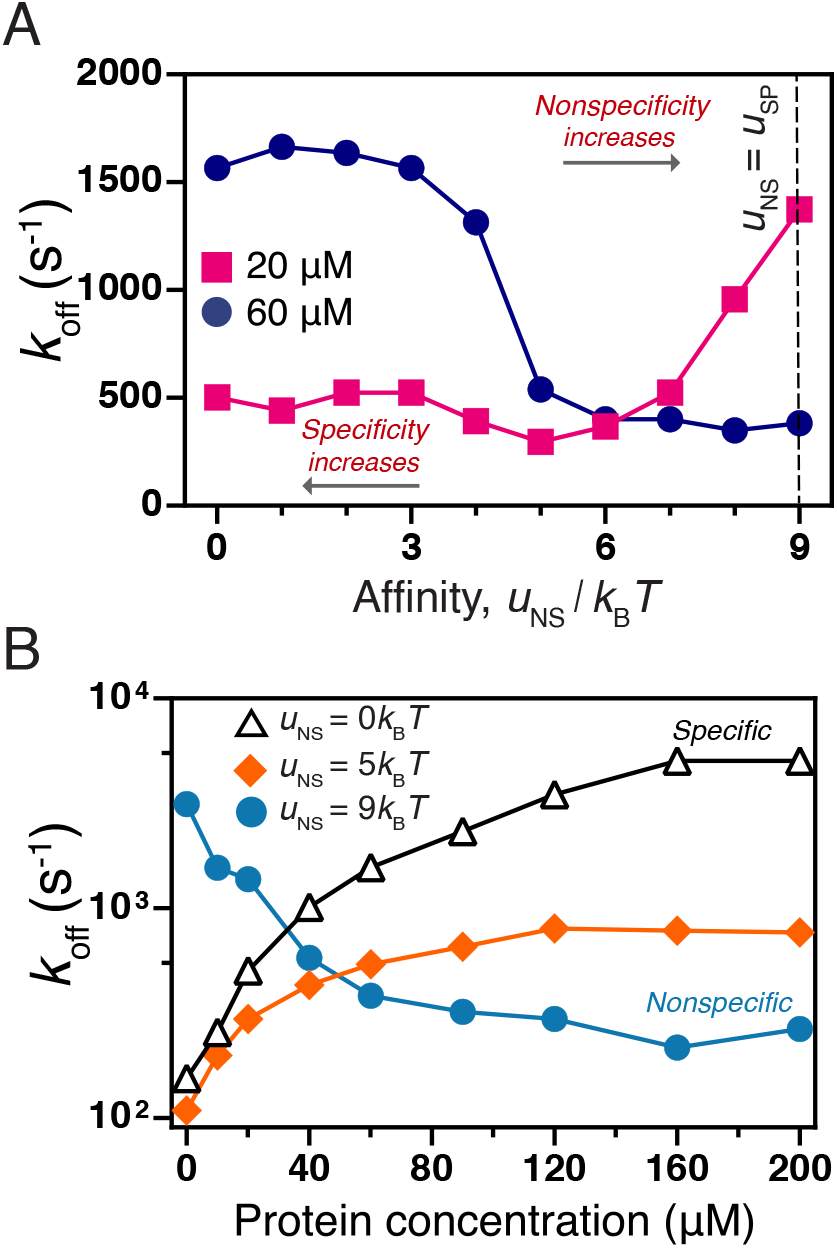
The role of nonspecificity on the facilitated dissociation of proteins. A) Off rates as a function of nonspecific protein-DNA attraction for two initially unbound protein concentrations of 20 and 60 *µ*M. The vertical line refers to non-specifically interacting proteins. B) The calculated off rates as a function of the initially unbound protein concentration. The proteins can bind to DNA via specific and nonspecific interactions (i.e., *u*_SP_ > 0 and *u*_NS_ > 0) or bind to DNA only specifically (i.e., *u*_SP_ > 0 and *u*_NS_ = 0). The specific attraction is *u*_SP_ = 9*k*_B_*T* in all cases. The data points are joined to guide the eye.

As the proteins distinguish less between specific and nonspecific DNA (i.e., *u*_NS_ → *u*_SP_), the dependence of the off-rate on the concentration is completely reversed (Figure 3A): Unlike the conventional FD, in which off rates raise with increasing concentration, the off rates of nonspecific proteins become slower as the concentration increases. This inverse-FD effect is more evident when the off rates are plotted as a function of the protein concentration (Figure 3B). As the unbound protein concentration goes up, the off-rate of nonspecific proteins slows down and reaches a plateau, at which the dissociation becomes roughly 10 times slower than the SD case (circles in Figure 3B). Note that in the calculation of the off rates, we do not distinguish between 1D and 3D escape of the proteins from binding sites. Overall, our simulations suggest that nonspecifically and specifically interacting structural proteins can exhibit marginally different kinetic responses to the increasing concentration of solution-phase proteins. In the next sections, we discuss these findings further by considering the relationship between chromosomal structure and dissociation kinetics.

### D. High NAP concentration leads to chromosome compaction

Nonspecific interactions between NAPs and DNA are known to contribute to chromosome organization [28, 51]. Thus, we suspect that as the proteins condense on the DNA polymer, the resulting structural alterations could affect the facilitated dissociation of the DNA-bound proteins by the solution-phase proteins. In order to quantify the chromosome compaction, we analyze the chromosome organization both visually and by calculating the quasiequilibrium radius of gyration of the DNA polymer, *R*_g_, for various nonspecific attractions and protein concentrations.

In our simulations, the initial configuration of the nucleiod structure fills roughly 40% of the overall cellular volume (Figure 1A). Depending on the protein levels and protein-DNA nonspecific attraction, the nucleiod either swells to fill the entire cell volume or collapses down to its smallest size, which is dictated by the steric interactions (Figure 4A-D). In a protein-free system or if the protein concentration is low (i.e., 1 copy per 500 bp or less), the size of the chromosome is mainly determined by the dimensions of the cellular confinement. For our chromosome model, this number is *R*_g_ ∼ 170 nm (the vertical line in Figure 4A and upper panels of Figure 4B, C and D). Notably, this open chromosome structure is in accord with the Hi-C contact maps of Fis deficient cells [52, 53]. We first consider the case, where proteins can bind the whole DNA but cannot form stable complexes with nonspecific DNA due to the low protein-DNA nonspecific attraction (in our simulations, this attraction limit is *u*_NS_ ≤ 3*k*_B_*T*). In this cases, the chromosome size has no apparent dependence on the protein concentration, and *R*_g_ is close to its protein-free value (Figure 4A, B). However, even in this swollen chromosomal structure, the proteins form clusters by bridging multiple specific (high affinity) binding sites (left panel of Figure 4E). Since, in our model, there is no protein-protein or DNA-DNA attractions, such DNA bridges are only possible if proteins are shared by multiple specific sites [34, 54]. Although the bridges anchor multiple protein-free DNA segments, they are unable to compact the chromosome even at the highest protein levels that we use here (i.e., 200 *µ*M or 1 protein per 60 bp). Hence, the resulting swollen polymer structure pervades the entire cellular volume regardless of the protein level (Figure 4B). Notably, the protein clusters are dynamic and contain on average several tens of proteins, and the average cluster size is not affected by the protein concentration (Supplementary Figure S4).

**FIG. 4.**
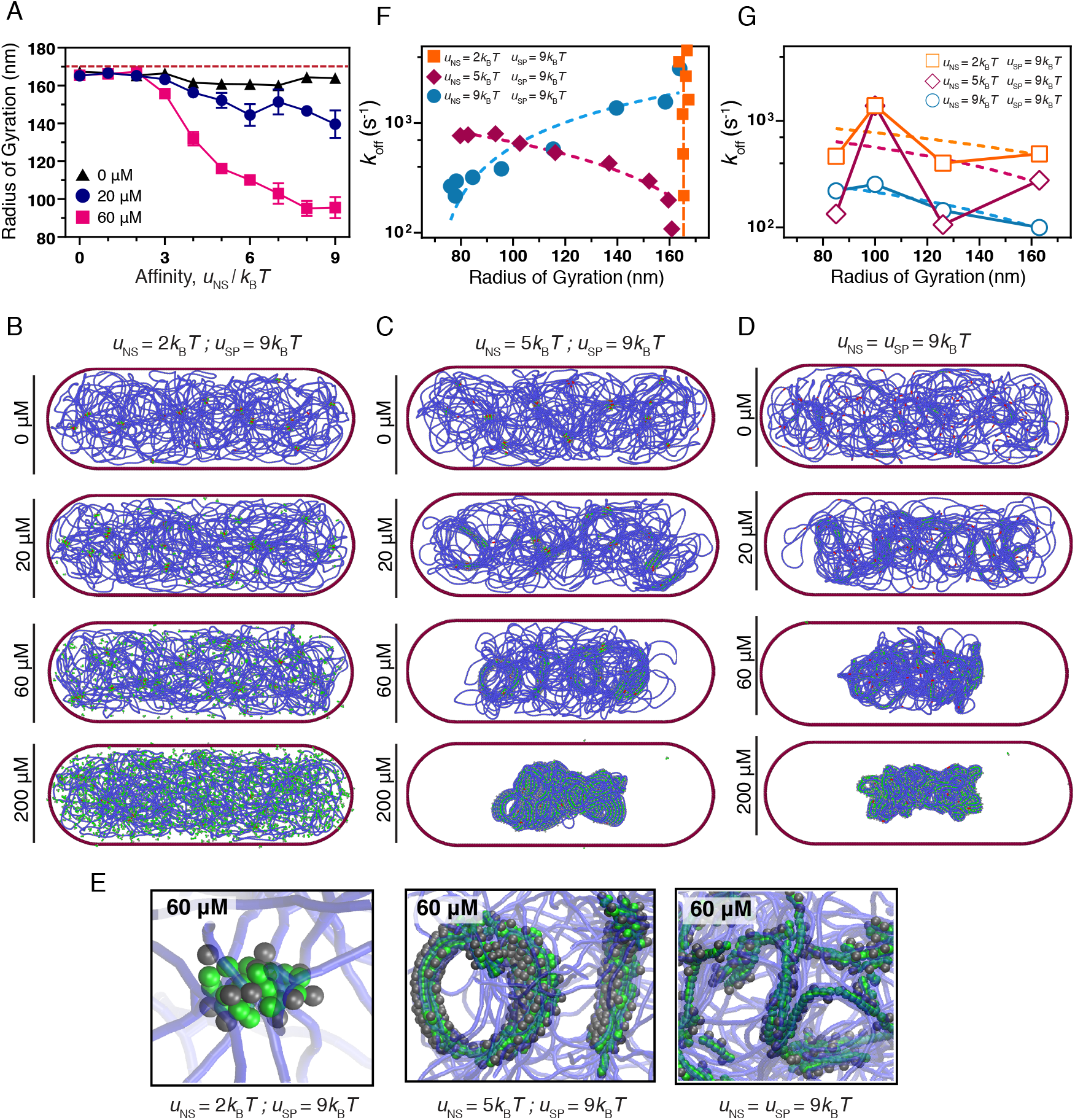
Chromosome compaction and facilitated dissociation. A) The calculated values of radius of gyration, *R*_g_, of chromosome structure as a function of nonspecific attraction for three concentrations of initially unbound proteins. The vertical line indicates the protein-free nucleoid size in the simulations. B, C, D) Representative snapshots of chromosomal structures for the initially unbound Fis concentrations of 0, 20, 60, and 200 *µ*M (left-to-right) for a specific (SP) binding energy of *u*_SP_ = 9*k*_B_*T*. Initially bound and unbound proteins are both marked by the same colors in the snapshots. The beads of the cell boundary are indicated by magenta. E) Close-up views of protein clusters for various concentrations for the cases in B,C,D. F) The off rates versus *R*_g_ for various protein-DNA cases. G) Off-rates obtained in a restrained DNA-polymer system as a function of pre-set *R*_g_ values. In F and G, the dashed lines are linear regression fits to guide the eye.

Contrarily, if proteins can form stable complexes with both specific and nonspecific DNA, chromosome compaction exhibits a highly concentration-dependent behaviour (Figure 4A-D). As the protein concentration is increased (i.e., higher than 20 *µ*M or more than 1 protein per ∼ 400 bp), the proteins form large (i.e., 40-50 proteins) and inter-connected multiprotein-DNA complexes (Figure 4C-E and Supplementary Figure S4). For specific proteins, we observe in our simulations that genomic regions containing relatively high-affinity binding sites act as nucleation sites for the formation of larger clusters. As the protein concentration is increased, the chromosomal structure shrinks in a concentration-dependent manner (Figure 4A-D). Notably, even though there is no torsional component in the DNA-polymer model, circular structures, in which multiple DNA segments are juxtaposed by specifically interacting proteins, emerge (Figure 4E). Nonspecific proteins, on the other hand, tend to form more 1D-like structures as compared to the specific ones as shown in Figure 4E in accord with the previous studies [34].

At the highest protein level that we consider here (i.e., 1 protein per 60 bp), the chromosome completely collapses into a globular structure similar to a polymer chain in poor solvent conditions (bottom panels in Figures 4C,D). Such collapse requires a nonspecific attraction greater than *u*_NS_ > 3*k*_B_*T*) (Figures 4A). Notably, nonspecific proteins seem to be more effective in compacting the chromosomal structure (Figures 4A). These protein-induced structures are consistent with the high resolution force-spectroscopy images of Fis-induced compaction of isolated DNA segments at similar protein levels [51]. Overall, our simulations show that nonspecific interactions significantly increase the capability of structural NAPs to compact the chromosomal structure in a concentration-dependent manner.

### E. Chromosome compaction affects protein-dissociation kinetics

Returning to our discussion on dissociation kinetics, in our calculations, the average off rates are determined upon the first dissociation events of the initially DNA-bound proteins. In some cases, the average time scales of chromosomal organization and protein dissociation are not well separated (i.e., a quasi-equilibrium polymer configuration is not reached at the instant of initial dissociation) (Supplementary Figure S5). However, a comparison between the off rates and chromosome size could serve to identify the mechanistic of the different concentration responses of nonspecific and specific DNA-binding proteins since nonspecificity affects the chromosomal compaction drastically (Figure 4A-E).

In Figure 4F, we compile the off rates of various protein-DNA interaction scenarios from Figures 2 and 3 and plot them as a function of the corresponding quasiequilibrium chromosome size, *R*_g_. The inverse concentration response of the off rates of specifically and nonspecifically DNA-binding proteins shown in Figure 3B also persists in Figure 4F. For specific proteins, the fastest off rates correspond to the minimum size of the chromosomal structure (at the maximum protein level). Contrarily, for nonspecific proteins, the fastest off rates correspond to the most open chromosomal structure (at the minimum protein level) (Figure 4F). If the proteins bind to nonspecific DNA very weakly (while forming stable complexes with a limited number of specific DNA sites), their off rates do not exhibit any change with *R*_g_ as expected (the orange data in Figure 4F).

As long as nonspecific attraction is strong enough, high NAP levels decreases the chromosome size and increase the nucleic acid concentration within the pervaded volume of the chromosome (Figure 4A). This compaction may cause a DNA-segmental type of FD [20] or change the frequency of protein rebinding events to the DNA polymer [55]. Hence, in order to decouple the compaction (or expansion) of the polymer from the protein dissociation, we restrain the self-avoiding walk configuration of the DNA polymer at prescribed *R*_g_ values (i.e., no thermal fluctuations for the polymer) and run a set of protein-dissociation simulations (Figure 4G). In these simulations, the proteins unbind from and bind to the DNA polymer but are unable to change its conformation. When we choose an unbound protein level of 60 *µ*M, for which we observe a considerable amount of FD-related effects (Figures 2 and 3), the calculated off rates show a systematic decrease as *R*_g_ is increased for all protein types (Figure 4G). Particularly for the nonspecific proteins, the compact chromosomal structure does not correlate with slow off rates and follow a dissociation pattern similar to the specific proteins (Figure 4G). Thus, based on our simulations with the restrained DNA polymers, we conclude that the chromosome compaction in general acts to increase the off rates of the DNA-bound proteins but nonspecific interactions can cause a reverse response (Figure 4F,G). Overall, our MD simulations suggest that chromosome compaction can affect the off rates of specific and nonspecific proteins in opposite ways.

### F. DNA-bound proteins are replaced by both lower and higher affinity proteins via FD

So far in our simulations, the initially DNA-bound and unbound proteins have equal binding attraction towards the DNA. Nevertheless, various NAPs and their mutants can coexist and compete for the same binding site with relatively small differences in their affinities [56, 57]. For instance, dissociation constants of K94A mutant and wild type Fis differ by a factor of ∼ 4 [56], which may result in ∼ 1*k*_B_*T* difference in the DNA-protein attraction (see Supplementary data for calculations). Moreover, DNA-bound Fis/HU was shown to exchange with HU/Fis molecules from solution in single-molecule experiments, indicating the heterotypic nature of FD [6]. Similarly, H-NS were shown to be displaced by *Salmonella enterica* regulator protein SsrB [33].

To reveal the competition between DNA-bound and free proteins from solution but with relatively lower/higher affinities, we perform additional simulations by setting an attraction difference of ∼ 1*k*_*B*_*T* between the initially unbound and initially DNA-bound proteins. For simplicity, we choose *u*_SP_ = 9*k*_B_*T* and *u*_NS_ = 2*k*_B_*T* for initially DNA-bound proteins, for which the chromosome compaction has no effect on the dissociation kinetics (Figure 4F). Our simulations show that both relatively lower (*u*_SP_ = 8*k*_B_*T*) and higher (*u*_SP_ = 10*k*_B_*T*) affinity proteins are able to accelerate the dissociation of the DNA-bound proteins efficiently (Figure 5A). However, the solution-phase proteins are more effective in stripping the lower affinity DNA-bound proteins off the DNA: Solution proteins with *u*_SP_ = 10*k*_B_*T* can facilitate the off-rate of *u*_SP_ = 9*k*_B_*T*-proteins ∼ 7 times more as compared to the initially free *u*_SP_ = 8*k*_B_*T*-proteins (Figure 5B). This suggests that even a small difference in affinities can play a key role in hierarchical dissociation of the proteins on a DNA.

**FIG. 5.**
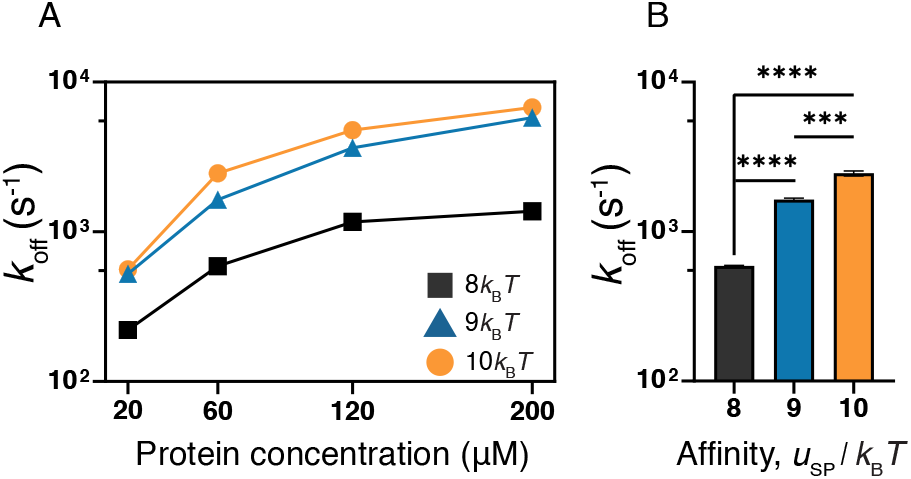
Heterotypic behaviour of FD. A) The off rates of proteins that interact with DNA with a specific binding strength of *u*_SP_ = 9*k*_B_*T* for various concentration of initially unbound proteins. The specific DNA binding attraction of the initially unbound proteins are *u*_SP_ = 8, 9, 10*k*_B_*T*. In all cases, the nonspecific attraction is *u*_NS_ = 2*k*_B_*T*. B) The off rates at a concentration of 60 *µ*M for three cases; when the initially unbound proteins have lower, equal, and higher affinity than initially DNA-bound proteins (unpaired, Two-Tailed, Student’s t test).

## IV. DISCUSSION

To summarize, our extensive MD simulations, where the DNA-bound proteins are allowed to unbind from a self-avoiding, circular, and confined DNA polymer, show that both specific and nonspecific NAPs modulate their off rates depending on their cellular concentration levels. As the concentration of solution-phase (DNA-free) structural proteins increases, the protein off rate increases for specifically DNA-binding proteins but decreases for nonspecific proteins (Figures 2 and 3). Further, mapping the calculated off-rate values to the corresponding chromosome size for each protein concentration reveals that specific proteins tend to stay shorter on their DNA recognition sites, whereas nonspecific proteins tend to stay longer in compact chromosomal structures (Figure 4F). Further, the simulations, where DNA-bound and solution-phase proteins have different affinity, demonstrate that the affinity determines the winner of the competition between the two different protein species for the same binding site, and the proteins with higher affinity can shorten the residence time of lower-affinity DNA-bound proteins (Figure 5). Our simulations also show that nonspecific DNA-protein attraction has a substantial effect on the off rates of proteins dissociating from a folded chromosomal structure even though the proteins unbind from their specific binding sites (Figure 2B). In the following sections, we will discuss our findings in the context of DNA-residence times and gene regulation in more detail.

### A. Concentration-dependent residence times of NAPs can have regulatory effects on transcription

As bacterial DNA binding proteins, TFs demonstrate sequence-specific recognition and binding behaviors. They control the flow of the genetic information via promoting or repressing transcription by enhancing the recruitment of RNA polymerase or preventing its binding to target DNA. First step to this process is the binding of a TF to its target site. When the cellular concentration of the TF increases, the probability of the protein to find its DNA target site increases linearly with the concentration [58]. Once the TF binds to its recognition site, up or down-regulation of the corresponding genes occurs. However, this view implicitly assumes that, *i*) the concentration only affects on rates (i.e., association rate) but have no effect on the off rates, which is defined as the reciprocal of the residence time of the protein on its DNA target site; and *ii*) once the TF is bound to its target site, how long it stays there does not change its transcriptional output.

A series of *in vitro* single-molecule studies have already challenged the first assumption and show that off rates of initially DNA-bound Gfp-Fis or HU becomes faster as the solution-phase concentration of the proteins was increased [6, 10, 11]. Our simulations go beyond those *in vitro* single-molecule experiments/simulations and demonstrate that structural dual-purpose proteins can exhibit concentration-dependent dissociation patterns also in a nucleoid-like environment, in which protein levels are directly coupled to chromosome organization. Our findings suggest that sequence specific proteins such as Fis can stay shorter on their target sites with increasing Fis levels, and thus, may couple their transcriptional activity with the duration of DNA occupancy. Proteins with no sequence specificity such as HU, on the other hand, can maintain their binding positions longer as their cellular levels rise. This property can allow them to form stable higher-level protein complexes and limit the gene accessibility effectively while facilitating the disassembly of the complexes as the protein levels fall (Figure 3). This view is also consistent with long DNA-occupancy times of HU and H-NS proteins (99%), which can assemble into large protein clusters on a DNA [59].

The second assumption to challenge is whether the transcription output of a gene depends on the residence time of the corresponding regulator or not. The competition ChIP experiments on the *Saccharomyces cerevisiae* Rap1 TF showed a direct correlation between the DNA residence time and transcriptional output: While transcriptional activation was coupled to long residence times, the low transcriptional output is correlated with rapid unbinding. [4]. Another *Saccharomyces cerevisiae* TF, Abf1, has also exhibited one order of magnitude variation in the off rates on different binding sites, each of which is related to a different functionality [5]. More recently, by tuning the number of nucleotide-recognizing repeat domains of transcription activator-like effectors (TALEs), a residence time-dependent decrease in the transcription levels of the glucocorticoid receptor-activated gene SGK1 was observed [3]. Thus, based on those findings and our results, we can postulate that if the residence time of a TF affects its transcription output, any physiological input modulating the residence times, such as the concentration of binding competitors, can also be a regulatory component. In the cases of dual-purpose proteins (i.e., Fis), this modulation is intermingled with alterations in chromosome structure induced by the protein itself.

### B. A tentative molecular picture for NAP-induced chromosome compaction

The high-resolution contact maps of *E. coli* chromosomes revealed a correlation between the high expression levels and chromosome structures that are rich in short-range contacts [53]. Furthermore, several fluorescence studies demonstrated that Fis, along with HN-S and RNA polymerase, can form foci in bacterial cells growing rapidly [60, 61]. The 3D STORM studies demonstrate that while Fis and HU can localize in large interconnected protein complexes (i.e., > 200 nm), H-NS can oligomarize into discrete nodes anchoring distant loci [62]. Our simulations may present a molecular picture for the formation of such potentially transcriptionally active protein-DNA structures in a concentration-dependent manner.

At low protein concentrations (i.e., ∼ 10 *µ*M), site-specific NAPs mainly manage to bridge multiple specific DNA binding sites into protein clusters (Figure 4E). These clusters are spread evenly throughout the relatively open bacterial chromosomes, consistent with super-resolution fluorescence microscopy [62]. Further, multiscale loop-like structures that are poor in high-affinity sites are anchored by these protein clusters (Figure 4B), confirming that such high-affinity sites can be functional in organizing DNA [54, 63]. Noteworthily, in our quasi-equilibrium configurations, the nonspecific DNA segments are mostly protein-free except those neighbouring the specific sites, suggesting the role of specific sites in the initiation of protein-DNA complexation from high-affinity sequences [54, 64].

As the concentrations of structural NAPs approach their *in vivo* peak levels (i.e., ∼ 50 *µ*M for Fis or ∼ 40 *µ*M for HU), two distinct behaviours emerge depending on the binding mode of the protein. For specifically interacting Fis-like proteins, the protein clusters become fewer upon merging, indicating the higher order organization of specific DNA binding sites (Figure 4C). For nonspecifically interacting proteins (e.g., HU), protein-DNA complexes tend to form elongated structures bridging multiple DNA segments (Figure 4D,E). We should underline that these protein-DNA assembles emerge despite the absence of a net attraction between the proteins and induce an attraction between remote DNA segments as observed for H-NS [34]. At excess protein levels of above ∼ 100 *µ*M, protein-DNA complexes shrink the chromosomes into highly compact structures (Figure 4C-E). All proteins are trapped inside or on the surface of such structure. Further, our simulations also suggest that nonspecifically interacting proteins seem to compact the chromosome more efficiently (Figure 4A) even though they tend to form filamentous protein-DNA structures. Such compact DNA structures can act like steric filters and may exclude the large-size macromolecules (i.e., ribosome) from the nucleoid [62].

For proteins that interact with DNA mostly specifically (e.g., their diffusion coefficients in bulk and on nonspecific DNA are similar), our simulations suggest a strong conventional FD but no protein-induced compaction (Figure 4B). Interestingly, the lack of H-NS were shown to cause minimal changes in the nucleoid volume as well [40], suggesting that such nonspecific NAPs may undergo FD at least in a loose chromosomal structure. Nevertheless, the compaction of chromosome by other structural NAPs can also change the DNA-residence times of these proteins in a chromosome-organization dependent way [31].

### C. Chromosome organization can affect the facilitated-dissociation response of DNA-bound proteins

The 3D chromosome structure is another component of transcriptional regulation [65–67]. Chromosome compaction or loosening can affect the recruitment of RNA polymerase. Our simulations also suggest that FD and protein-induced chromosomal alterations could function together to modulate the gene accessibility by changing DNA-residence times (Figure 4F). Our MD simulations at prescribed chromosome-compaction levels also show that in general, compaction promotes a quicker dissociation in accord with DNA-segmental facilitate dissociation (Figure 4G). However, this architectural regulation can vary for specific and nonspecific NAPs differently as we demonstrate here (Figure 4). Additionally, a TF protein may respond differently to the concentration fluctuations depending on DNA sequences surrounding a promoter site. Accordingly, apo and holo forms of metal-sensing

TFs show chromosome-compaction-dependent residence times but oppositely in live *E. coli* cells [31, 32].

### D. FD, chromosome organization and the functional diversity of Fis

Fis can participate in the regulation of more than 200 genes including itself either as an activator or repressor necessary for growth, replication, and energy metabolism [23, 56, 68–70]. Despite the high coarsegraining level of our dimeric protein model, our simulations could offer some clues about the broad functional diversity of Fis, as follows: *i*) During the early stages of the growth, a gradual increase in Fis levels can modulate Fis residence times differently on different target sequences via facilitated dissociation, *ii)* As Fis concentration increases, its negative autoregulation capacity may decrease via FD (i.e., it cannot exclude RNA polymerase effectively from the target sites due to fast turnover rates). Particularly, through the lag phase to the early exponential phase, this behaviour can pursuit, and *iii)* Fis could also lead to a temporal DNA compaction via protein-dense clusters at the regions containing Fis regulated genes. Depending on the affinity of nonspecific sites surrounding the gene or physical proximity of other specific sites nearby the gene, DNA-compaction may inhibit RNA polymerase’s access to these gene-rich regions in a selective way,

To see if there is a correlation between Fis concentration and transcriptional activity, we revisit the time courses of cellular Fis and transcript levels under nutritional upshift from the literature (see Figure 2 of Ref [23]). Since the expression of Fis is significantly regulated at the transcriptional level, protein abundance is expected to be similar to the transcript levels with a time lag of ∼ 10 minutes ([23]. Accordingly, when the cellular level of Fis reaches halfway through its maximum, the transcript level of Fis is already at its maximum. Consistently, the available DNA microarray expression profiles show a cumulative upregulation pattern for Fis-regulated genes within the early exponential phase [69]. Once the Fis level attains its maximum at around >50 *µ*M (i.e., the level at which we observe high DNA compaction and dense protein-DNA arrays in our simulations), Fis levels begin to decrease [23, 69]. The decrease continues from the early logarithmic phase to the stationary phase. Within this interval, the transcript levels of Fis also decreases from its peak level to vanishing values, suggesting a decreasing e regulatory activity of Fis. We speculate that this reduction could be due to DNA compaction and/or dense-protein clusters that can extend the residence times of Fis on DNA and thus its respressive activity.

In conclusion, our extensive MD simulations suggest that off rates of proteins falling off from a folded DNA polymer could be correlated with cellular protein levels and the chromosomal organizational changes that the proteins cause. Cumulatively, these effects could regulate the protein’s biomolecular functionality by modulating its genome-wide residence times. An investigation of genome-wide off rates for proteins such Fis by using dynamic ChIP or other experimental tools [4, 5] would reveal such mechanisms. Finally, beyond prokaryotic cells, the phenomenon that we demonstrate here is very likely to be functional in eukaryotic cell nuclei as well, given that some structural DNA-binding proteins, such as histone proteins, have already been shown to undergo facilitated dissociation [15, 71].

## ACKNOWLEDGEMENTS

AE acknowledges J.F. Marko for the discussion on the Fis binding energies and Dr. B. Hekimoglu and Dr. E.J. Banigan for their careful readings of the manuscript. AGA is supported by TUBITAK 2247-C STAR program. This research is supported by TUBITAK 2232 (2018/4) 119C010.

## Data availability

All the Python codes can be accessed at https://github.com/ZaferKosar/FD.git

## References

[1] Douglas F. Browning and Stephen J. W. Busby. The regulation of bacterial transcription initiation. Nature Reviews Microbiology, 2:57–65, January 2004.

[2] Aswin Sai Narain Seshasayee, Karthikeyan Sivaraman, and Nicholas M. Luscombe. An Overview of Prokaryotic Transcription Factors, pages 7–23. Springer Netherlands, Dordrecht, 2011.

[3] Karen Clauß, Achim P Popp, Lena Schulze, Johannes Hettich, Matthias Reisser, Laura Escoter Torres, N Henriette Uhlenhaut, and J Christof M Gebhardt. DNA residence time is a regulatory factor of transcription repression. Nucleic Acids Research, 45(19):11121–11130, 2017.

[4] Colin R Lickwar, Florian Mueller, Sean E Hanlon, James G McNally, and Jason D Lieb. Genome-wide protein-DNA binding dynamics suggest a molecular clutch for transcription factor function. Nature, 484(7393):1–7, 2012.

[5] Wim J de Jonge, Mariël Brok, Philip Lijnzaad, Patrick Kemmeren, and Frank CP Holstege. Genome-wide off-rates reveal how DNA binding dynamics shape transcription factor function. Molecular Systems Biology, 16(10):1405–16, 2020.

[6] J S Graham, Reid C Johnson, and J F Marko. Concentration-dependent exchange accelerates turnover of proteins bound to double-stranded DNA. Nucleic Acids Research, 39(6):2249–2259, 2011.

[7] Joseph J Loparo, Arkadiusz W Kulczyk, Charles C Richardson, and Antoine M van Oijen. Simultaneous single-molecule measurements of phage T7 replisome composition and function reveal the mechanism of polymerase exchange. Proceedings of the National Academy of Sciences, 108(9):3584–3589, 2011.

[8] Chandra P Joshi, Debashis Panda, Danya J Martell, Nesha May Andoy, Tai-Yen Chen, Ahmed Gaballa, John D Helmann, and Peng Chen. Direct substitution and assisted dissociation pathways for turning off transcription by a MerR-family metalloregulator. Proceedings of the National Academy of Sciences, 109(38):15121–15126, 2012.

[9] Simone Kunzelmann, Caroline Morris, Alap P Chavda, John F Eccleston, and Martin R Webb. Mechanism of Interaction between Single-Stranded DNA Binding Protein and DNA. Biochemistry, 49(5):843–852, 2010.

[10] Ramsey I Kamar, Edward J Banigan, Aykut Erbaş, Rebecca D Giuntoli, Monica Olvera de la Cruz, Reid C Johnson, and John F Marko. Facilitated dissociation of transcription factors from single DNA binding sites. Proceedings of the National Academy of Sciences, 114(16):E3251–E3257, 2017.

[11] N Hadizadeh, Reid C Johnson, and J F Marko. Facilitated dissociation of a nucleoid protein from the bacterial chromosome. Journal of bacteriology, 2016.

[12] Tai-Yen Chen, Yu-Shan Cheng, Pei-San Huang, and Peng Chen. Facilitated Unbinding via Multivalency-Enabled Ternary Complexes: New Paradigm for Protein-DNA Interactions. Accounts of Chemical Research, 51(4):860–868, 2018.

[13] S R Joseph, M Palfy, L Hilbert, M Kumar, J Karschau Elife, and 2017. Competition between histone and transcription factor binding regulates the onset of transcription in zebrafish embryos. elifesciences.org.

[14] Tina Lebar, Anamp x0017E e Verbiamp x0010D, Ajasja Ljubetiampx0010D, and Roman Jerala. Polarized displacement by transcription activator-like effectors for regulatory circuits. Nature Chemical Biology, pages 1–13, 2018.

[15] Sinan Kilic, Andreas L Bachmann, Louise C Bryan, and Beat Fierz. Multivalency governs HP1. Nature Communications, 6:1–11, 2015.

[16] Y Luo, J A North, S D Rose, and M G Poirier. Nucleosomes accelerate transcription factor dissociation. Nucleic Acids Research, 42(5):3017–3027, 2014.

[17] Bryan Gibb, Ling F Ye, Stephanie C Gergoudis, YoungHo Kwon, Hengyao Niu, Patrick Sung, and Eric C Greene. Concentration-Dependent Exchange of Replication Protein A on Single-Stranded DNA Revealed by Single-Molecule Imaging. PLOS ONE, 9(2):e87922, 2014.

[18] Charles E Sing, Monica Olvera de la Cruz, and John F Marko. Multiple-binding-site mechanism explains concentration-dependent unbinding rates of DNA-binding proteins. Nucleic Acids Research, 42(6):3783–3791, 2014.

[19] Christoffer Åberg, Karl E Duderstadt, and Antoine M van Oijen. Stability versus exchange: a paradox in DNA replication. Nucleic Acids Research, 44(10):4846–4854, 2016.

[20] Rebecca D Giuntoli, Nora B Linzer, Edward J Banigan, Charles E Sing, Monica Olvera de la Cruz, John S Graham, Reid C Johnson, and John F Marko. DNA-Segment-Facilitated Dissociation of Fis and NHP6A from DNA Detected via Single-Molecule Mechanical Response. Journal of molecular biology, 427(19):3123–3136, 2015.

[21] Aykut Erbaş, Monica Olvera de la Cruz, and John F Marko. Receptor-Ligand Rebinding Kinetics in Confinement. Biophysical Journal, 116(9):1609–1624, 2019.

[22] T Ali Azam, A Iwata, A Nishimura, S Ueda, and A Ishihama. Growth phase-dependent variation in protein composition of the Escherichia coli nucleoid. Journal of bacteriology, 181(20):6361–6370, 1999.

[23] C A Ball, R Osuna, K C Ferguson, and Reid C Johnson. Dramatic changes in Fis levels upon nutrient upshift in Escherichia coli. Journal of bacteriology, 174(24):8043–8056, 1992.

[24] Charles J. Dorman and Kelly A. Kane. DNA bridging and antibridging: a role for bacterial nucleoid-associated proteins in regulating the expression of laterally acquired genes. FEMS Microbiology Reviews, 33(3):587–592, 05 2009.

[25] Arlene Kelly, Martin D Goldberg, Ronan K Carroll, Vittoria Danino, Jay C D Hinton, and Charles J Dorman. A global role for Fis in the transcriptional control of metabolism and type III secretion in Salmonella enterica serovar Typhimurium. Microbiology, 150(Pt 7):2037–2053, 2004.

[26] Zhong Qian Sankar L Adhya Subhash C Verma. Architecture of the Escherichia coli nucleoid. pages 1–35, December 2019.

[27] John van Noort, Sander Verbrugge, Nora Goosen, Cees Dekker, and Remus Thei Dame. Dual architectural roles of HU: formation of flexible hinges and rigid filaments. Proceedings of the National Academy of Sciences of the United States of America, 101(18):6969–6974, 2004.

[28] Dunja Skoko, Daniel Yoo, Hua Bai, Bernhard Schnurr, Jie Yan, Sarah M McLeod, John F Marko, and Reid C Johnson. Mechanism of Chromosome Compaction and Looping by the Escherichia coli Nucleoid Protein Fis. Journal of molecular biology, 364(4):777–798, 2006.

[29] Dunja Skoko, Ben Wong, Reid C Johnson, and John F Marko. Micromechanical Analysis of the Binding of DNA-Bending Proteins HMGB1, NHP6A, and HU Reveals Their Ability To Form Highly Stable DNAProtein Complexes †. Biochemistry, 43(43):13867–13874, 2004.

[30] Soumya G Remesh, Subhash C Verma, Jian-Hua Chen, Axel A Ekman, Carolyn A Larabell, Sankar Adhya, and Michal Hammel. Nucleoid remodeling during environmental adaptation is regulated by HU-dependent DNA bundling. Nature Communications, pages 1–12, 2020.

[31] Tai-Yen Chen, Ace George Santiago, Won Jung, L ukasz Krzemiński, Feng Yang, Danya J Martell, John D Helmann, and Peng Chen. Concentration-and chromosomeorganization-dependent regulator unbinding from DNA for transcription regulation in living cells. Nature Communications, 6:7445, 2015.

[32] Won Jung, Kushal Sengupta, Brian M Wendel, John D Helmann, and Peng Chen. NAR Breakthrough Article Biphasic unbinding of a metalloregulator from DNA for transcription (de)repression in Live Bacteria. pages 1–10, 2020.

[33] Don Walthers, You Li, Yingjie Liu, Ganesh Anand, Jie Yan, and Linda J Kenney. Salmonella enterica Response Regulator SsrB Relieves H-NS Silencing by Displacing H-NS Bound in Polymerization Mode and Directly Activates Transcription. The Journal of biological chemistry, 286(3):1895–1902, 2011.

[34] C A Brackley, S Taylor, A Papantonis, P R Cook, and D Marenduzzo. Nonspecific bridging-induced attraction drives clustering of DNA-binding proteins and genome organization. Proceedings of the National Academy of Sciences of the United States of America, 110(38):E3605–E3611, 2013.

[35] Suckjoon Jun and Bela Mulder. Entropy-driven spatial organization of highly confined polymers: lessons for the bacterial chromosome. Proceedings of the National Academy of Sciences of the United States of America, 103(33):12388–12393, 2006.

[36] J Dorier and A Stasiak. Modelling of crowded polymers elucidate effects of double-strand breaks in topological domains of bacterial chromosomes. Nucleic Acids Research, 41(14):6808–6815, 2013.

[37] Kurt Kremer and Gary S Grest. Dynamics of entangled linear polymer melts: A molecular-dynamics simulation. Journal of Chemical Physics, 92(8):5057–5086, 1990.

[38] Gary S Grest and Michael Murat. Structure of grafted polymeric brushes in solvents of varying quality: a molecular dynamics study. Macromolecules, 26(12):3108–3117, 1993.

[39] Stephen P Hancock, Stefano Stella, Duilio Cascio, and Reid C Johnson. DNA Sequence Determinants Controlling Affinity, Stability and Shape of DNA Complexes Bound by the Nucleoid Protein Fis. PLOS ONE, 11(3):e0150189, 2016.

[40] Yunfeng Gao, Yong Hwee Foo, Ricksen S Winardhi, Qingnan Tang, Jie Yan, and Linda J Kenney. Charged residues in the H-NS linker drive DNA binding and gene silencing in single cells. Proceedings of the National Academy of Sciences of the United States of America, 114(47):12560–12565, 2017.

[41] Priyanka Gawade, Gaurav Gunjal, Anamika Sharma, and Payel Ghosh. Reconstruction of transcriptional regulatory networks of fis and h-ns in escherichia coli from genome-wide data analysis. Genomics, 112(2):1264–1272, 2020.

[42] Aykut Erbaş, Monica Olvera de la Cruz, and John F Marko. Effects of electrostatic interactions on ligand dissociation kinetics. Physical Review E, 97(2-1):022405, 2018.

[43] Min-Yeh Tsai, Bin Zhang, Weihua Zheng, and Peter G Wolynes. Molecular Mechanism of Facilitated Dissociation of Fis Protein from DNA. Journal of the American Chemical Society, 2016.

[44] S Cocco, J F Marko, and R Monasson. Stochastic Ratchet Mechanisms for Replacement of Proteins Bound to DNA. Physical Review Letters, 112(23):238101–5, 2014.

[45] Katelyn Dahlke and Charles E Sing. Facilitated Dissociation Kinetics of Dimeric Nucleoid-Associated Proteins Follow a Universal Curve. Biophysical Journal, 112(3):543–551, 2017.

[46] Steve Plimpton. Fast parallel algorithms for short-range molecular dynamics. Journal of Computational Physics, 117(1):1–19, 1995.

[47] Charles R. Harris, K. Jarrod Millman, Stfan J. van der Walt, Ralf Gommers, Pauli Virtanen, David Cournapeau, Eric Wieser, Julian Taylor, Sebastian Berg, Nathaniel J. Smith, Robert Kern, Matti Picus, Stephan Hoyer, Marten H. van Kerkwijk, Matthew Brett, Allan Haldane, Jaime Fernndez del Ro, Mark Wiebe, Pearu Peterson, Pierre Grard-Marchant, Kevin Sheppard, Tyler Reddy, Warren Weckesser, Hameer Abbasi, Christoph Gohlke, and Travis E. Oliphant. Array programming with NumPy. Nature, 585(7825):357–362, September 2020.

[48] W Humphrey, A Dalke, and K Schulten. VMD: visual molecular dynamics. Journal of molecular graphics, 14(1):33–38, 1996.

[49] Alexander Stukowski. Visualization and analysis of atomistic simulation data with OVITO-the Open Visualization Tool. MODELLING AND SIMULATION IN MATERIALS SCIENCE AND ENGINEERING, 18(1), JAN 2010.

[50] Yongping Shao, Leah S Feldman-Cohen, and Robert Osuna. Functional Characterization of the Escherichia coli Fis–DNA Binding Sequence. Journal of molecular biology, 376(3):771–785, February 2008.

[51] R Schneider, R Lurz, G Lüder, C Tolksdorf, A Travers, and G Muskhelishvili. An architectural role of the Escherichia coli chromatin protein FIS in organising DNA. Nucleic Acids Research, 29(24):5107–5114, 2001.

[52] Ankit Gupta, Abdul Wasim, and Jagannath Mondal. Nucleoid associated proteins and their effect on E. coli chromosome. bioRxiv, page 2020.11.05.369934, 2020.

[53] Virginia S Lioy, Axel Cournac, Martial Marbouty, Stéphane Duigou, Julien Mozziconacci, Olivier Espéli, Frédéric Boccard, and Romain Koszul. Multiscale Structuring of the E. coli Chromosome by Nucleoid-Associated and Condensin Proteins. Cell, 172(4):771–783.e18, 2018.

[54] Jian Zhang, Yvonne Zeuner, Alexandra Kleefeld, Gottfried Unden, and Andreas Janshoff. Multiple sitespecific binding of Fis protein to Escherichia coli nuoA-N promoter DNA and its impact on DNA topology visualised by means of scanning force microscopy. Chembiochem : a European journal of chemical biology, 5(9):1286–1289, 2004.

[55] Azita Parsaeian, Monica Olvera de la Cruz, and John F Marko. Binding-rebinding dynamics of proteins interacting nonspecifically with a long DNA molecule. Physical Review E, 88(4):675, 2013.

[56] Leah S Feldman-Cohen, Yongping Shao, Derrick Meinhold, Charmi Miller, Wilfredo Colon, and Robert Osuna. Common and Variable Contributions of Fis Residues to High-Affinity Binding at Different DNA Sequences. Journal of bacteriology, 188(6):2081–2095, 2006.

[57] S P Hancock, T Ghane, D Cascio, R Rohs, R Di Felice, and Reid C Johnson. Control of DNA minor groove width and Fis protein binding by the purine 2-amino group. Nucleic Acids Research, 41(13):6750–6760, 2013.

[58] S E Halford and J F Marko. How do site-specific DNA-binding proteins find their targets? Nucleic Acids Research, 32(10):3040–3052, 2004.

[59] Mathew Stracy, Jakob Schweizer, David J Sherratt, Achillefs N Kapanidis, Stephan Uphoff, and Christian Lesterlin. Transient non-specific DNA binding dominates the target search of bacterial DNA-binding proteins. Molecular Cell, 81(7):1499–1514.e6, 2021.

[60] T A Azam, S Hiraga, and A Ishihama. Two types of localization of the DNA-binding proteins within the Escherichia coli nucleoid. Genes to cells : devoted to molecular & cellular mechanisms, 5(8):613–626, 2000.

[61] Wenqin Wang, Gene-Wei Li, Chongyi Chen, X Sunney Xie, and Xiaowei Zhuang. Chromosome Organization by a Nucleoid-Associated Protein in Live Bacteria. Science, 333(6048):1445–1449, 2011.

[62] Wenqin Wang, Gene-Wei Li, Chongyi Chen, X Sunney Xie, and Xiaowei Zhuang. Chromosome Organization by a Nucleoid-Associated Protein in Live Bacteria. Science, 333(6048):1445–1449, 2011.

[63] Aleksandre Japaridze, Sylvain Renevey, Patrick Sobetzko, Liubov Stoliar, William Nasser, Giovanni Dietler, and Georgi Muskhelishvili. Spatial organization of DNA sequences directs the assembly of bacterial chromatin by a nucleoid-associated protein. The Journal of biological chemistry, 292(18):7607–7618, 2017.

[64] M Bétermier, D J Galas, and M Chandler. Interaction of Fis protein with DNA: bending and specificity of binding. Biochimie, 76(10-11):958–967, 1994.

[65] Ümit Pul and Rolf Wagner. Nucleoid-Associated Proteins: Structural Properties, pages 149–173. Springer Netherlands, Dordrecht, 2010.

[66] Charles J Dorman. Genome architecture and global gene regulation in bacteria: making progress towards a unified model? pages 1–7, April 2013.

[67] Douglas F Browning, David C Grainger, and Stephen J W Busby. Effects of nucleoid-associated proteins on bacterial chromosome structure and gene expression. Current Opinion in Microbiology, 13(6):773–780, 2010.

[68] W Ross, J F Thompson, J T Newlands, and R L Gourse. E.coli Fis protein activates ribosomal RNA transcription in vitro and in vivo. The EMBO Journal, 9(11):3733–3742, 1990.

[69] Meranda D Bradley, Michael B Beach, A P Jason de Koning, Timothy S Pratt, and Robert Osuna. Effects of Fis on Escherichia coli gene expression during different growth stages. Microbiology, 153(9):2922–2940, 2007.

[70] Christina Kahramanoglou, Aswin S. N. Seshasayee, Ana I. Prieto, David Ibberson, Sabine Schmidt, Jurgen Zimmermann, Vladimir Benes, Gillian M. Fraser, and Nicholas M. Luscombe. Direct and indirect effects of H-NS and Fis on global gene expression control in Escherichia coli. Nucleic Acids Research, 39(6):2073–2091, 11 2010.

[71] D T Gruszka, S Xie, H Kimura, and H Yardimci. Single-molecule imaging reveals control of parental histone recycling by free histones during DNA replication. Science advances, 6(38), 2020.

