## Supplemental Data and Figures for "Facilitated Dissociation of Nucleoid Associated Proteins from DNA in the Bacterial Confinement"

### Supplementary Data and Figures for Facilitated Dissociation of Nucleoid Associated Proteins from DNA in the Bacterial Confinement

Zafer Koşar[1], A. Göktuĝ Attar[1,2], and Aykut Erbaş[1]

[1] *UNAM-National Nanotechnology Research Center and Institute of Materials Science & Nanotechnology, Bilkent University, Ankara 06800, Turkey and*

[2] *Department of Molecular Biology & Genetics, Bilkent University, Ankara 06800, Turkey*

**Details of Molecular Dynamics (MD) simulations and data analyses**

**Simulation Model**

In our MD simulations, a circular DNA molecule and proteins are modeled using the coarse-grained Kremer-Grest (KG) bead-spring chains in implicit solvent. The nucleic acid chain is a smaller version of the full-size *E. coli* genome with $1.2 \times 10^5$ base pairs (bp), which is modelled with $N = 12000$ KG beads with each bead representing $\sim 10$ bp. The circular DNA chain and proteins are confined by a rigid concentric cylinder with an aspect ratio of 3. The caps of the cylinders are built as semi-spheres. The volume of the confinement is adjusted in accordance with the nucleic acid concentration of *E. coli*, which corresponds to a DNA volume fraction of $\sim 1\%$.

The simulation systems consist of two types of proteins: initially DNA-bound and free proteins. Initially, bound proteins are positioned near promoter regions (red beads represent the binding sites that are spaced evenly along the DNA polymer. Given the smaller size of our model genome, the number of binding sites is chosen as $n_0 = 120$, which is lower than the consensus sequences determined but high enough to perform statistical analyses of unbinding kinetics. To monitor the concentration-dependent dissociation, a prescribed concentration of initially free proteins, $c_{\mathrm{f}}$, are added at random positions in the simulation boxes in addition to the $n_0 = 120$ initially bound proteins. The total protein concentration in cells ranges between 12 $\mu$M (only DNA-bound proteins) and 212 $\mu$M, which correspond to $n_0 = 120$ and $n_0 = 2225$ protein copies per cell, respectively.

DNA polymer, proteins, and the confinement boundary are composed of identical beads of size $b = 1\sigma$, where $\sigma$ is the unit length in the MD simulations. The steric interactions between these beads are modeled by a truncated and shifted Lennard-Jones (LJ) potential, also known as WCA,

$$V^{\mathrm{LJ}}(r) = \begin{cases} 4u\left[(\sigma/r)^{12} - (\sigma/r)^6 + v_s\right] & r \leq r_c \\ 0 & r > r_c. \end{cases} \tag{1}$$

The cut-off distance $r_c = 2^{1/6}\ \sigma$, the shift factor $v_s = 1/4$, and the interaction strength $u = 1\ k_{\mathrm{B}}T$ are chosen to obtain good solvent conditions (i.e., a repulsive interaction) unless otherwise noted. Here, $k_{\mathrm{B}}$ is the Boltzmann constant, and $T$ is the absolute temperature.

For the attractive interactions between DNA and proteins, we set $r_c = 2.5\ \sigma$ and $v_s = 0$. The interaction strength value is set to $u \gg 1$ to model nonspecific and specific interactions (see below for details). There is no net attraction between the beads of the proteins. The bonding between the adjacent beads of the DNA chain or proteins is taken care of by a nonlinear FENE potential

$$V^{\rm Bond}(r) = -0.5kr_0^2 \ln\left[1 - (r/r_0)^2\right], \tag{2}$$

where the default value of the bond stiffness is $k = 30\ k_{\rm B}T/\sigma^2$. In Eq. 2, the distance between adjacent beads is $r = |\mathbf{r}|$, and the maximum bond length is $r_0 = 1.5\ \sigma$. In order to main the average "cherry" shape of the dimeric protein, a harmonic potential with reference angle of $\Theta_0 = 50^\circ$ and stiffness of $k_\Theta = 12k_{\rm B}T/rad^2$ is also set between the three beads.

A harmonic bending potential is introduced to model the semi-flexible nature of the DNA as

$$V^{\rm Bend}(\Theta) = k_\Theta\left(\Theta - \Theta_0\right)^2, \tag{3}$$

where the potential strength is $k_\Theta = 15k_{\rm B}T/rad^2$, $\Theta$ is the angle formed by three adjacent beads, and $\Theta_0$ is the reference angle and is $\Theta_0 = \pi$ is for the DNA chain. These parameters provide an average persistence length of $\sim$15 beads, which is in accord with the persistence length of naked double-stranded DNA (i.e., $\sim$150 bp $\sim$ 50 nm).

All MD simulations are run with LAMMPS MD package at constant volume $V$ and reduced temperature $T_{\rm r} = 1.0$. In the preparation of the setups, each DNA polymer is prepared as a perfect circle of $N$ beads and is larger than the dimensions of the cellular confinement. In order to fit the polymer within the designated volume, the polymer is collapsed by assigning temporary self-attraction energy of 17 $k_{\rm B}T$ between all chain beads. Upon transferring the collapsed polymer inside the cylinders, this attraction is reversed to the default values, and then, the polymer is allowed to relax for $> 2 \times 10^4$ MD steps. Then, the initially bound proteins are placed on the positions of the respective binding sites. To check if the relaxation of the long polymers has any effect on the results, a shorter chain with $N = 2400$ is tested, but no quantitative difference in unbinding kinetics is observed (Figure S1). The data production runs are carried out for $10^6 - 10^9$ MD steps depending on the binding strength and protein concentration. During the simulations, DNA stays within the cellular volume, and occasionally a few proteins (among hundreds) leave the cylinder.

The simulations are run with a time step of $\Delta t = 0.005\ \tau$, where the unit time scale in the simulations is $\tau$, which is roughly the self-diffusion time of a single bead in dilute solution. The monomeric LJ mass is $m = 1$ for all beads. The temperature is kept constant by a Langevin thermostat with a thermostat coefficient $\gamma = 0.5\ \tau^{-1}$. The volume of the total simulation box is set to $L_x \times L_y \times L_z = 184 \times 64 \times 64\ \sigma^3$. Periodic boundary conditions are used in all directions.

##### Estimation of relative strengths of non/specific protein-DNA interactions

In order to estimate the relative difference in binding energies between the specific (SP) and nonspecific (NS) interactions, we use the dissociation constants reported in the literature According, if $K_\mathrm{d}$ values for SP and NS differ roughly a factor of $\sim 1000$, the difference in the binding attraction could be obtained by using

$$K_\mathrm{d}^\mathrm{SP}/K_\mathrm{d}^\mathrm{NS} \sim \ln\left(\Delta u/k_\mathrm{B}T\right), \tag{4}$$

as $\Delta u \sim 7k_\mathrm{B}T$. The same approximation was also used to estimate the difference between the binding energies of various Fis mutants. T Note that in Eq. 4, we assume that entropic contributions to the non/specific unbinding free energies are similar. In reality, the specific binding can cause higher entropy difference due to relatively more stable binding of the protein to the recognition site as compared to the nonspecific binding.

##### Rescaling of the simulation-time unit to the real-time unit

We map the diffusion time of a molecule of size $d = 6.8$ nm in simulation units to the real time units by using the Einstein diffusion equation

$$\tau = \frac{d^2}{6D}, \tag{5}$$

where $D$ is the diffusion of the particle and can be estimated to be $D = 0.5k_BT/2\pi\eta d$. Here the viscosity of water is $\eta = 10^{-3}$ N.s/m$^2$, $k_BT \approx 4 \times 10^{-21}$ Nm. Here we also use a prefactor of 0.5 to account for the higher water viscosity in the crowded cell environment.

If we combine all of our equation, we obtain

$$\tau = \frac{\pi \eta d^3}{k_B T}, \tag{6}$$

which leads to $\tau \approx 250$ ns. We use this conversion to map all of our calculated off rates to the units of 1/s and simulation times to the units of milliseconds.

##### Calculation of protein-cluster sizes and distributions

In order to quantify both the number of clusters and protein population in each cluster, we post-process simulation trajectories . There are several clustering algorithms available for clustering problems. For instance, a widespread clustering algorithm is k-NN (k-Nearest Neighbor), widely used in machine learning. Although this algorithm applies to a wide range of practices, it requires the cluster number as an input. In our case, the quantities of the protein clusters vary drastically with different conditions, even among the replicas of the same conditions. Therefore, it would not be practical to use such an algorithm. Our next option was to use a density-based algorithm.

Here, the best suiting approach was to use Stillinger's definition of the cluster. In his 1962 paper, Stillinger defines a cluster as units in the range of each other for a specified radius. Therefore, clusters can grow in any direction thus, forming various shapes and sizes. Investigating how many clusters there are and which protein belongs to which cluster requires a multistep strategy. First, we list all the neighbors for each protein. We immediately eliminated the TFs with no neighboring TF to reduce complexity, thus, the computation runtime. Then, each neighbor list is compared with the rest. The lists with common proteins are pooled together, establishing the clusters. The combined lists represent the clusters. Therefore, the number of elements in a combined list is the equivalent of cluster size. Also, the number of the lists gives the cluster count in a system. We decided on a threshold as 5 proteins (i.e., the smallest cluster size). Consequently, any protein complex with less than 5 proteins is not considered a cluster. Conventionally, the threshold distance for a protein to be counted as a neighbour is 1.5 times the bead diameter. However, after benchmarking, we found it more suitable to use 2.1 times the size of the beads (i.e., $2.1\sigma$, where $\sigma$ is the simulation unit length scale) following the visual inspection and a quantitative

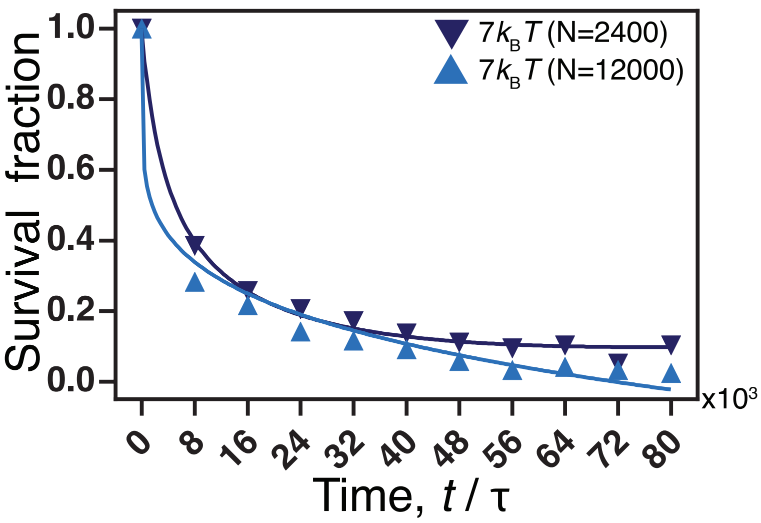


FIG. S1. Comparison of survival fractions of small ($N = 2400$) and large ($N = 12000$) systems at $u_{SP} = 7k_BT$ in the absence of free TF Protein. Here, nonspecific interaction (NS) energy is set to $u_{NS} = 1.6k_BT$ in both cases.

quantification of the cluster sizes. For instance, when the entire polymer collapses into a globular structure at our maximum protein concentration of 200 $\mu$M, we expect to a single super cluster including all the proteins in the cell volume. Starting with $1.6\sigma$, and gradually increase by $0.1\sigma$ led to one single cluster at $2.1\sigma$. Expanding the threshold radius joined other proteins to the clusters, which are essentially not part of the multiprotein complex. Also, decreasing the radius $2.1\sigma$ even by $0.05\sigma$ excluded the beads that belong to the complex. The cluster population and sizes are shown in Figure SS4 for various cases.

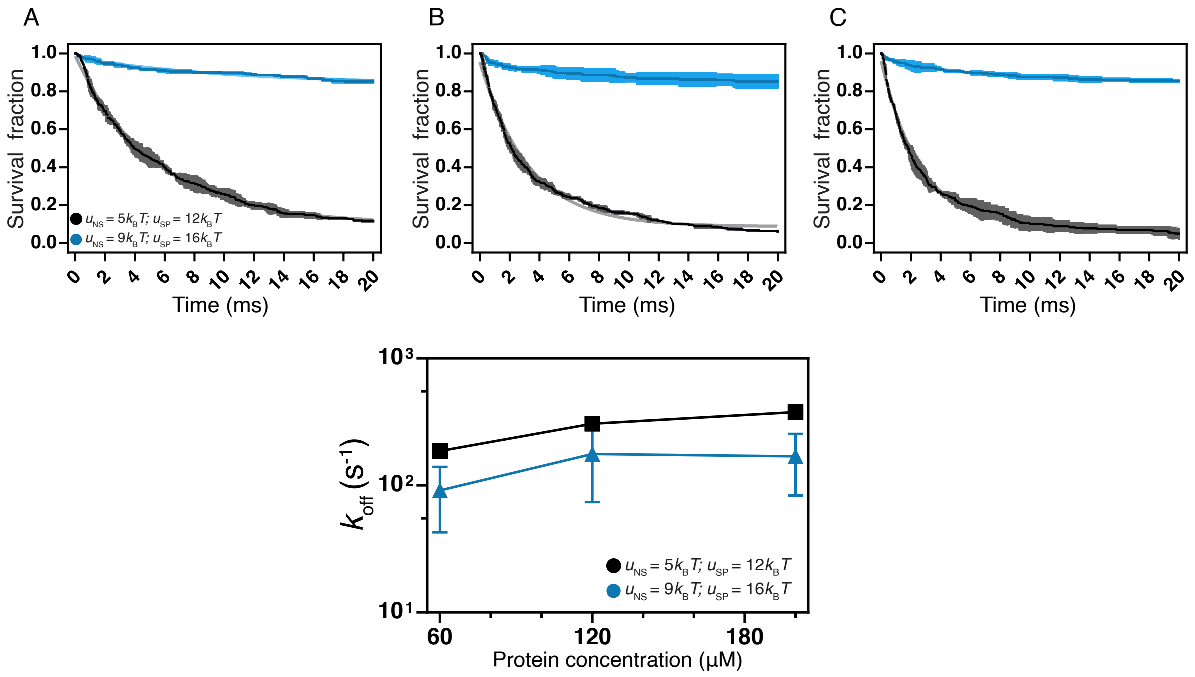


FIG. S2. Comparison of various specific and nonspecific binding energies. Upper panel shows the survival fractions for 0 (blue) and 60 $\mu$ M (black) initially unbound proteins for two different attraction sets, each which is higher than the case used in the main text. Lower panel shows the of rates extracted from the survival fraction data.

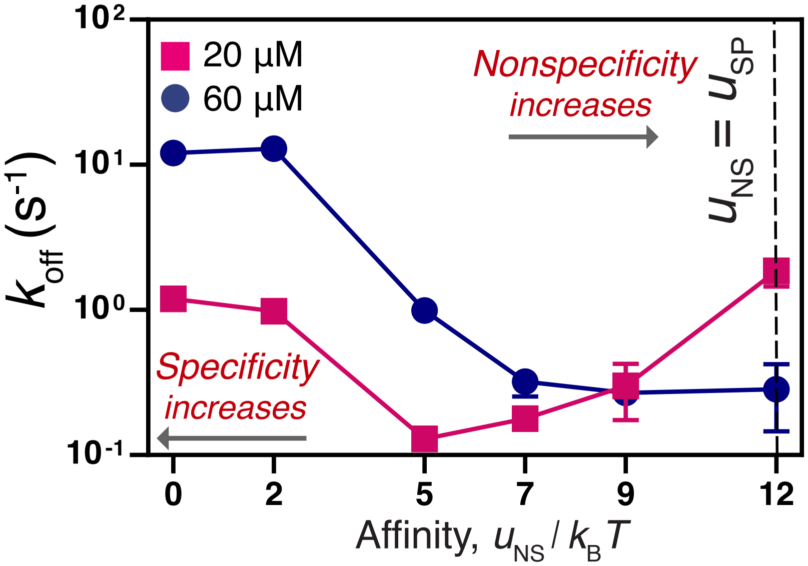


FIG. S3. Off-rates as a function of nonspecific protein-DNA attraction for two initially unbound protein concentrations of 20 and 60 $\mu$M. The vertical line refers to nonspecifically interacting proteins. The specific attraction is $u_{\rm SP} = 12k_{\rm B}T$ in all cases. The data points are joined to guide the eye. In the simulation both Langevin and NVT algorithms were applied.

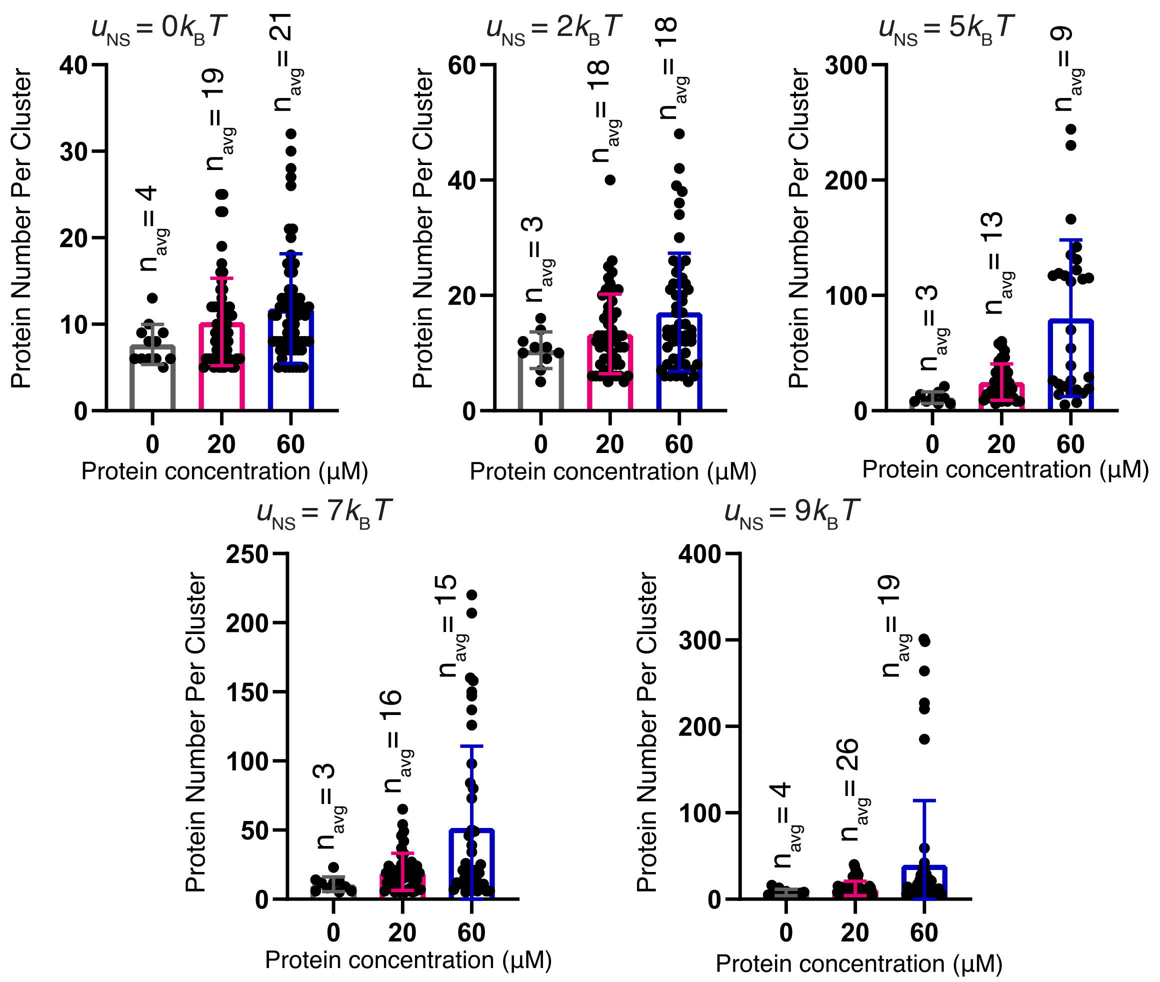


FIG. S4. Size of protein clusters for various nonspecific attraction values and for three protein concentrations. The specific attraction is $u_{\mathrm{SP}} = 9k_{\mathrm{B}}T$ in all cases. $n_{avg}$ refers to the average number of clusters.

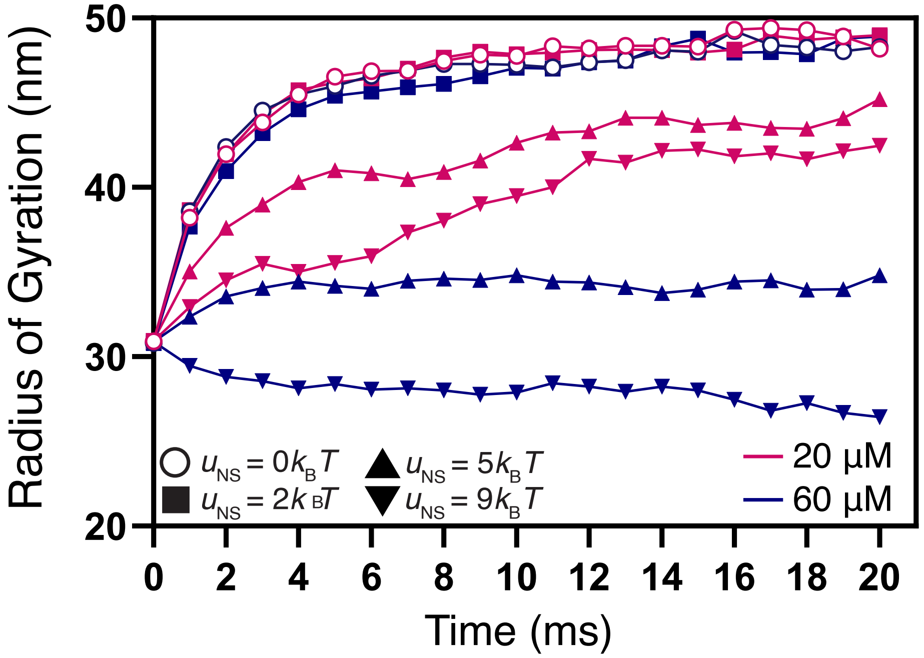


FIG. S5. Characteristic size of the chromosomes as a function of the time for two protein concentrations and nonspecific attractions. The specific attraction is $u_{\mathrm{SP}} = 12k_{\mathrm{B}}T$ in all cases.
